# Differential Control of Inhibitory and Excitatory Nerve Terminal Function by Mitochondria

**DOI:** 10.1101/2024.05.19.594864

**Authors:** Kirsten Bredvik, Timothy A. Ryan

## Abstract

Inhibitory neurons shape the brain’s computational landscape and rely on different cellular architectures and intrinsic properties than excitatory neurons. Maintenance of the overall balance of excitatory (E) versus inhibitory (I) drive is essential, as disruptions can lead to neuropathological conditions, including autism and epilepsy. Metabolic perturbations are a common driver of E/I imbalance but differential sensitivity of these two neuron types to metabolic lesions is not well understood. Here, we characterized differences in presynaptic bioenergetic regulation between excitatory and inhibitory nerve terminals using genetically encoded indicators expressed in primary dissociated neuronal cultures. Our experiments showed that inhibitory nerve terminals sustain higher ATP levels than excitatory nerve terminals arising from increased mitochondrial metabolism. Additionally, mitochondria in inhibitory neurons play a greater role in buffering presynaptic Ca^2+^ and inhibitory mitochondrial Ca^2+^ handling is differentially regulated by TMEM65-mediated acceleration of mitochondrial Ca^2+^ extrusion following bursts of activity. These experiments thus identify differential reliance on mitochondrial function across two major neuron types.

## Introduction

The central nervous system uses about 20% of the body’s energy, with neurons making up about 95% of the brain’s energy expenditure (Attwell & Laughlin, 2001). Previous work has shown that neuronal activity creates large fluctuations in the energetic burden of presynaptic terminals, requiring precise regulatory mechanisms to tightly control ATP production (Rangaraju et al., 2014; Pathak et al., 2015; Ashrafi et al., 2020). These previous investigations, however, did not attempt to address potential variations in these properties across different neuronal types. Several lines of evidence suggest that excitatory and inhibitory neurons may differ in aspects of metabolic control of their function, as metabolic perturbations in the brain can lead to excitation/inhibition (E/I) imbalances (Almannai et al., 2021; Tang & Monani, 2021).

Inhibitory neurons are known to regulate their firing rate, waveform shape, excitability, and development independently from excitatory neurons (Frank et al., 2001; Price et al., 2005; Bar et al., 2022, Mayer et al., 2018). A higher density of mitochondria has been observed in inhibitory neurons when compared to excitatory neurons (Kageyama & Wong-Riley, 1982; Nie & Wong-Riley, 1995; Gulyás et al., 2006), and *in vivo* studies have posited that functioning mitochondria are important in parvalbumin positive inhibitory neurons for sustaining high frequency firing rates and modulating murine sensory gating, socialization, and anxiety related behavior (Inan et al., 2016; Kontou et al., 2021). In recent years, single cell RNA sequencing and cell type specific proteomic analysis revealed intriguing differences in the expression of mitochondrial proteins between excitatory and inhibitory neuronal subtypes (Zeisel et al., 2015; Huntley et al., 2020; Wynne et al., 2021; Fecher et al., 2019). To understand how metabolic factors contribute to changes in circuit functioning in states such as neurodegeneration and psychiatric conditions (Li & Sheng, 2022), it is important to understand the differences in metabolism between excitatory and inhibitory synapses. Currently, a detailed exploration of the regulation and functioning of mitochondria in inhibitory neurons across metabolic conditions represents a fundamental knowledge gap in the field of neuronal metabolism.

To investigate possible metabolic differences in inhibitory versus excitatory neurons we took advantage of the recent identification of minimal enhancer elements (Mich et al., 2021; Dimidschstein et al., 2016) to drive the expression of a next-generation genetically-encoded ATP reporter, iATPSnFR2.0, mitochondrial targeted GCaMP6f and synaptophysin-pHluorin in inhibitory neurons and examined potential differences in metabolic control of nerve terminal functional of excitatory versus inhibitory neurons. Our data reveal that inhibitory axons have higher densities of mitochondria that results in much higher resting ATP levels than in excitatory axons. Furthermore, the increased mitochondrial content results in a larger total Ca^2+^ buffering capacity such that blocking mitochondrial Ca^2+^ uptake has a much bigger impact on nerve terminal Ca^2+^ accumulation, in turn impacting synaptic vesicle exocytosis. We additionally discovered that Ca^2+^ handling in inhibitory neuronal mitochondria was substantially different than in excitatory neuronal mitochondria, as inhibitory neuronal mitochondria relied on a putative NCLX regulator previously identified in muscle that accelerates mitochondrial Ca^2+^ extrusion (Teng et al., 2022). These results identify critical differences in the impact of mitochondrial function in inhibitory versus excitatory nerve terminals and will provide a useful framework in the future for how to understand potential metabolic drivers of disease states in the brain.

## Results

### Inhibitory axons have much higher mitochondrial basal ATP production than excitatory axons

To target inhibitory and excitatory neurons individually, we used the hDlxI56i enhancer and the Ca^2+^/calmodulin-dependent protein kinase II (CaMKII) promoter as expression control elements, respectively, that we previously showed to offer excellent specificity for selective expression in these two neuron classes in primary dissociated neuronal cultures (Farrell et al., 2024). The hDlxI56i construct contains a beta globin minimal promoter and the regulatory intergenic region found between the paired genes *Dlx5* and *Dlx6*, expressing across major transcriptomic subtypes of interneurons in the adult cortex (Mich et al., 2021; Dimidschstein et al., 2016).

Given that *in-vivo* inhibitory neurons are reported to have higher densities of mitochondria compared to excitatory neurons (Nie & Wong-Riley, 1995), we sought first to determine if this property is recapitulated in dissociated primary neuronal cultures. To characterize mitochondrial density in the excitatory and inhibitory neurons of our culture system, we co-expressed a mitochondrially targeted RFP (mito-RFP) along with an axonal marker (synaptophysin-pHluorin) in either hDlxI56i enhancer or CaMKII promoter expression constructs (from here on referred to as inhibitory and excitatory, respectively; Figure 1A). Axons were identified based on the expression of pHluorin and mitochondria were quantified in linearized images of the RFP channel (Figure 1B). In agreement with previous studies (Gulyás et al., 2006), we observed a ∼ 25% higher density of mitochondria per unit length of axon in inhibitory axons (∼ 1.5 per 10 µM axon) when compared to excitatory axons (∼ 1.2 per 10 µM axon).

**Figure 1.**
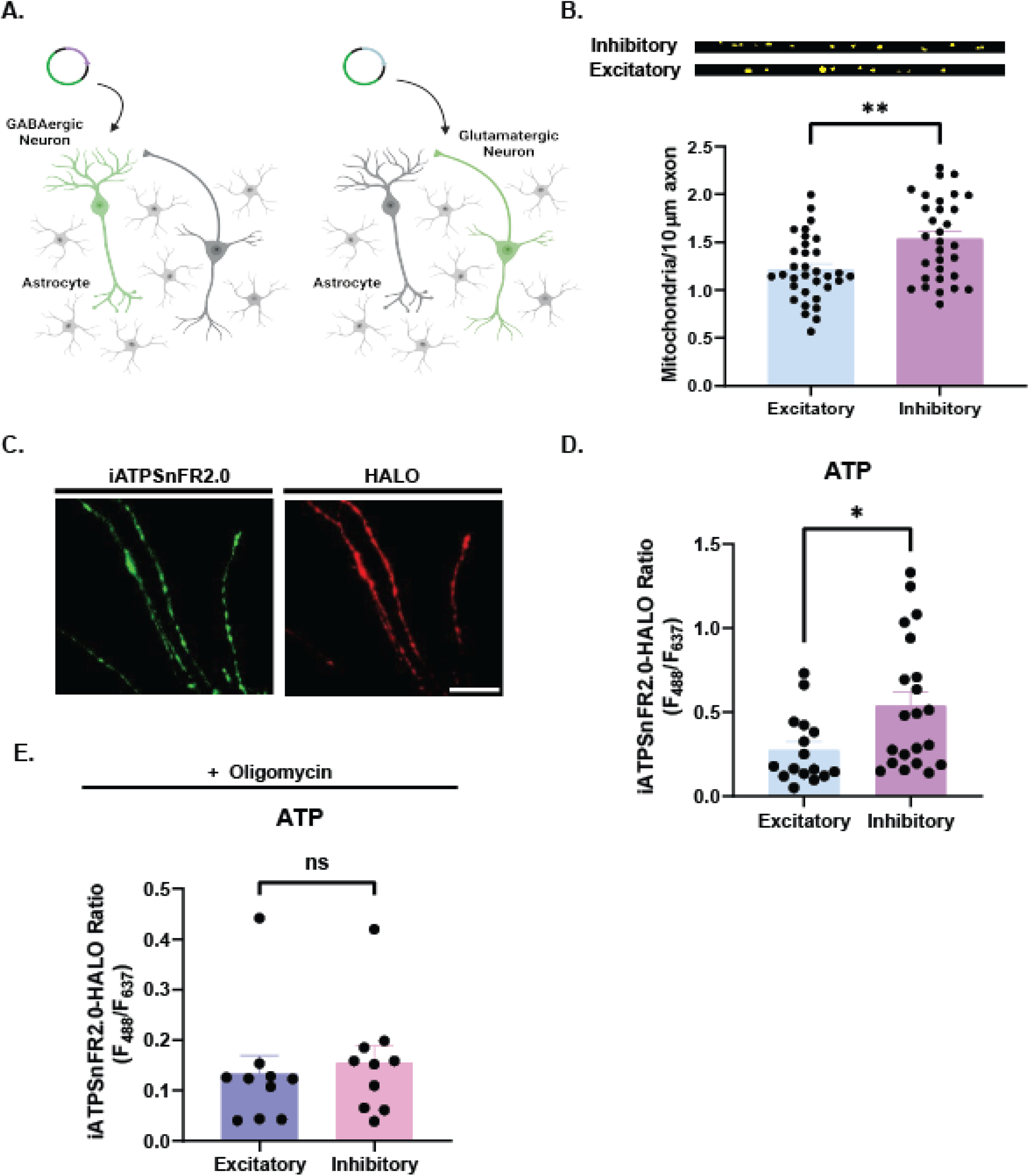
Resting ATP level differs between excitatory and inhibitory neurons. **A)** Schematic of cell type specific transfection of fluorescent reporter into GABAergic or glutamatergic neurons in culture. **B)** (Above) Representative images of linearized axon segments from excitatory and inhibitory neurons transfected with mitochondrially targeted RFP under CMV promoter (mito-RFP) and hDlxI56i (inhibitory) or CaMKII (excitatory) axon-targeted reporter (synaptophysin-pHluorin), respectively. (Below) Number of mitochondria per 10 µM axon length, quantified as peaks exceeding 110% of baseline fluorescence in linearized axon images (n=33 excitatory, mean=1.214. n=31 inhibitory, mean=1.536. Error bars SEM, p=0.0052, Mann-Whitney test). **C)** Representative images of cortical inhibitory neurons transfected with hDlxI56i iATPSnFR2.0-HALO, with 488 (left) and 637 (right) nm excitation (Scale bar 20 µM). **D)** Quantification of resting iATPSnFR2.0/HALO fluorescence ratio in cortical excitatory (CaMKII, blue) or inhibitory (hDlxI56i, purple) cells in 5 mM glucose (n=16 excitatory, mean=0.2742, n=21 inhibitory, mean=0.5379. *p=0.0123, Mann-Whitney test, error bars SEM). **E)** Quantification of resting iATPSnFRH2.0/HALO fluorescence ratio in excitatory and inhibitory neurons in presence of oligomycin with 1.1 mM glucose (n=10, excitatory mean=0.1322, inhibitory mean=0.1550. Error bars SEM, n.s., p=0.3527, Mann-Whitney test).

Based on the greater mitochondrial density of inhibitory neurons, we hypothesized that the metabolic state of inhibitory neurons might differ from that of excitatory neurons. To compare the metabolic state of excitatory and inhibitory neurons directly, we expressed the ratiometric ATP reporter iATPSnFR2.0-HALO (Marvin et al., 2024) in either inhibitory or excitatory neurons (Figure 1C, 1D) and quantified the ratio of the ATP sensitive channel (488 nm excitation) fluorescence to the ATP-insensitive (HALO + JF635/637 nm excitation) channel. These experiments revealed that inhibitory axons have almost 2-fold higherATP concentration than excitatory axons (Figure 1D). This difference in resting ATP however was eliminated with acute blockade of the F1-F0-ATP synthase (complex V) using oligomycin (Figure 1E), indicating that the large difference in resting ATP was primarily due to the increased respiratory capacity of inhibitory neurons. These data show that even though the mitochondrial density in inhibitory axons was only ∼ 25% greater than that of excitatory axons, it corresponded to an almost 2-fold change in resting ATP concentration.

### Oxidative fuels provide better metabolic support in inhibitory versus excitatory axons during bursts of electrical activity

We previously showed that excitatory nerve terminals rely on regulated ATP production to compensate for the energetic burdens associated with presynaptic function (Rangaraju et al., 2014; Ashrafi et al., 2017; Ashrafi et al., 2020; Marvin et al., 202). In response to a sustained burst of action potential (AP) firing, both excitatory and inhibitory axons showed a similar decrease in ATP levels during stimulation (Figure 2A, 2B; Supplementary Figure 1A-1C) that recovered to pre-stimulus levels over the next ∼ 75 s (Supplementary Figure 1A-1C) in the presence of glucose. When we eliminated the possible contributions of mitochondrial ATP production in both neuron types by blocking complex V with oligomycin, the temporal profile of ATP changes during and after a burst of AP firing were similar (Figure 2C, 2D). Restricting axons to purely glycolytic support did not lead to differences in the kinetics of ATP recovery following stimulation between these two types of neurons (Supplementary Figure 1D). Overall, these results indicated that glycolytic ATP production can maintain ATP homeostasis in both excitatory and inhibitory neuron types even with exhaustive stimulation, and therefore that both types have sufficient glycolytic capacity to sustain this type of activity.

**Figure 2.**
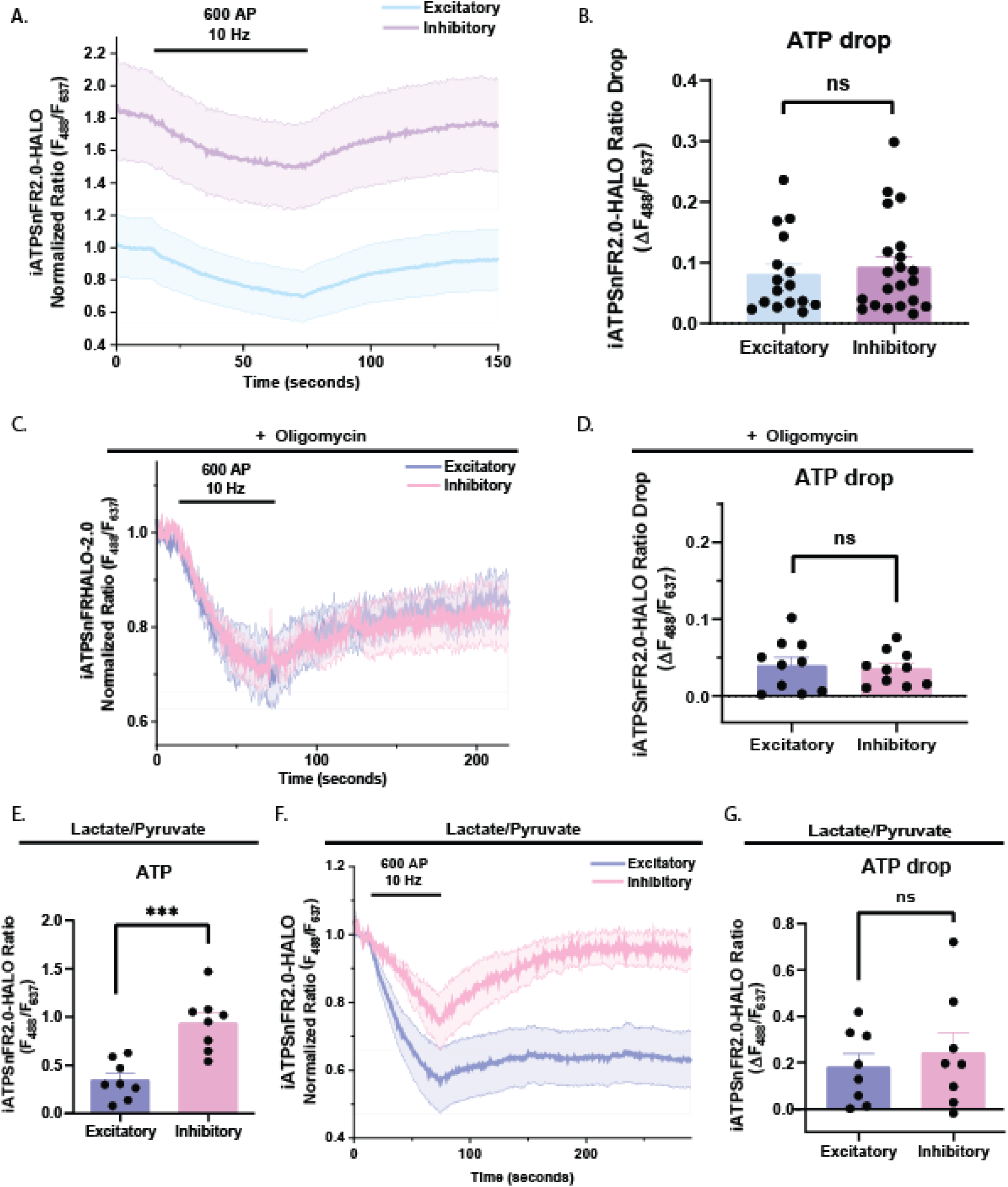
Inhibitory neurons fuel synaptic transmission with mitochondrial ATP more than excitatory neurons. **A)** iATPSnFR2.0/HALO fluorescence traces of excitatory and inhibitory neurons stimulated with 600 action potential (AP) at 10 Hz in 5 mM glucose, normalized to average initial fluorescence ratio of excitatory neurons prior to stimulation (n=16 excitatory, n=21 inhibitory, error bands SEM). **B)** Absolute drop in ATP ratio quantified at end of stimulation (n=16 excitatory, mean=0.081, n=21 inhibitory, mean=0.09296. Error bars SEM, n.s., p=0.7276, Mann-Whitney test). **C)** iATPSnFR2.0-HALO traces from excitatory and inhibitory neurons stimulated with 600 AP, 10 Hz during treatment with oligomycin in 1.1 mM glucose, normalized to ratio prior to stimulation (n=10 excitatory, n=10 inhibitory, error bands SEM). **D)** Quantification of absolute ATP drop following 600 AP, 10 Hz stimulation in 1.1 mM glucose and oligomycin, from traces in B) (n=10 excitatory, mean=0.03973. n=10 inhibitory, mean=0.03595. Error bars SEM, n.s., p=0. 9705, Mann-Whitney test). **E)** Quantification of resting iATPSnFRH2.0/HALO fluorescence ratio in excitatory and inhibitory neurons provided with 1.25 mM pyruvate and 1.25 mM lactate as fuel (n=8, excitatory mean=0.3477, inhibitory mean=0.9403. Error bars SEM, ***p=0.0006, Mann-Whitney test). **F)** iATPSnFR2.0-HALO traces of excitatory and inhibitory neurons stimulated with 600 AP at 10 Hz in lactate/pyruvate, normalized to ratio prior to stimulation (n=8, error bands SEM). **G)** Quantification of absolute ATP drop at end of stimulation in lactate/pyruvate, from traces in F) (n=8, excitatory mean=0.1897, inhibitory mean=0.2402. Error bars SEM, n.s., p>0.99, Mann-Whitney test).

*In-vivo* the source of fuel to sustain brain function likely varies depending on the brain region and various metabolic conditions. Although glucose is considered the primary fuel for the brain, certain neurons may rely on oxidative fuels such as lactate (Karagiannis et al., 2021), ketones (Chowdhury et al., 2014), or fatty acids (Kumar et al., 2023) under some conditions. To examine how ATP homeostasis might be impacted by forcing axons to rely solely on a mitochondrial fuel, we carried out ATP measurements under conditions where glucose was replaced by a mixture of lactate and pyruvate (1.25 mM each). Under these conditions the differences between resting ATP levels were ∼ 2.5-fold greater in inhibitory neurons compared to excitatory neurons (Figure 2E), consistent with our observation of higher mitochondrial densities in this neuron class (Figure 1B). In the presence of a purely respiratory fuel, sustained AP firing led to a similar drop in ATP levels in both neuron types (Figure 2F, 2G). The recovery of ATP levels following stimulation differed significantly in the two neuron populations, with inhibitory neurons revealing a more rapid and complete recovery following stimulation (Supplementary Figure 1E, 1F).

### Inhibitory nerve terminals have increased mitochondrial Ca^2+^ buffering

We previously showed that following increases in cytoplasmic Ca^2+^, axonal mitochondria in excitatory neurons rapidly take up Ca^2+^ from the cytoplasm during electrical activity through the mitochondrial Ca^2+^ uniporter (MCU) in a manner that is independent of efflux from the endoplasmic reticulum (ER; Ashrafi et al., 2020). In order to determine whether axonal mitochondria in inhibitory neurons are similarly sensitive to cytoplasmic Ca^2+^ and operate independently from ER Ca^2+^ fluxes, we expressed synaptically- and mitochondrially-targeted Ca^2+^ indicators (synaptophysin-GCaMP6f, Mito4x-jRCaMP1b) in inhibitory neurons and examined the mitochondrial Ca^2+^ signal in response to AP firing in the presence or absence of cyclopiazonic acid (CPA), a blocker of the ER ATP-dependent Ca^2+^ uptake pump (Supplementary Figure 2). These experiments showed that as with excitatory synapses (Supplementary Figure 2A, 2C-D) and axonal mitochondria (Supplementary Figure 2E-F), Ca^2+^ uptake into inhibitory synapses (Supplementary Figure 2B-D) and axonal mitochondria do not rely on Ca^2+^ efflux from the ER during electrical stimulation (Supplementary Figure 2E-F).

The greater density of mitochondria in inhibitory neurons suggests that mitochondrial Ca^2+^ uptake might more effectively buffer the changes in cytoplasmic Ca^2+^ than that of excitatory neuronal mitochondria. To test this hypothesis, we expressed an shRNA targeting MCU that was previously validated for use in our culture system (Ashrafi et al., 2020) in both excitatory and inhibitory neurons along with the synaptically-targeted Ca^2+^ indicator synaptophysin-GCaMP6f. In the absence of shRNA targeting MCU, Ca^2+^ dynamics in excitatory and inhibitory nerve terminals following a 20 AP burst of firing showed similar profiles (Figure 3A, 3B) including peak amplitudes, extrusion kinetics and resting pre-stimulus Ca^2+^ levels (Figure 3A-D). Similar to previous findings (Ashrafi et al., 2020), loss of MCU function had no significant impact on Ca^2+^ dynamics following a burst of AP firing in excitatory nerve terminals (Figure 3A). In contrast, loss of MCU in inhibitory neurons led to an almost 2-fold increase in nerve terminal cytoplasmic Ca^2+^ during the same stimulation (Figure 3B), strongly implying that in this neuronal class the increased mitochondrial density has a profound influence on synaptic function. To further test this hypothesis, we examined the impact of loss of MCU on exocytosis in both excitatory and inhibitory neuron types by co-expressing the MCU shRNA and the exocytic reporter synaptophysin-pHluorin (Sankaranarayanan et al., 2000). In excitatory nerve terminals, loss of MCU had no measurable impact on the amplitude of the pHluorin response following a 100 AP burst of firing (Figure 3E, 3G). In contrast, loss of MCU in inhibitory neurons led to a doubling of the exocytic peak during the same stimulus (Figure 3F, 3G). These results demonstrate that mitochondria have a more profound influence on presynaptic function in inhibitory compared to excitatory axons.

**Figure 3.**
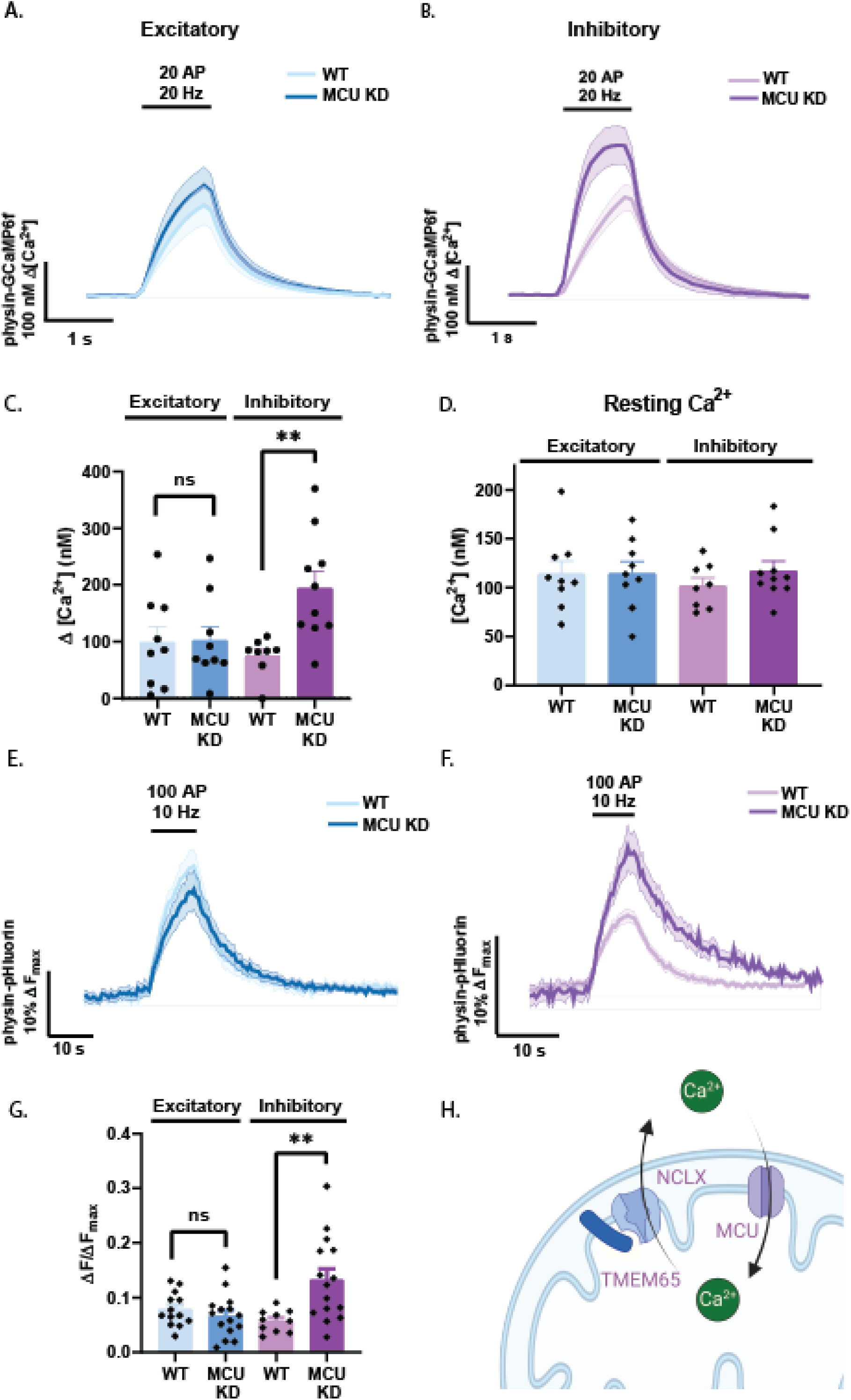
The mitochondrial calcium uniporter (MCU) is a pivotal regulator of synaptic transmission in inhibitory neurons. **A)** Synaptic Ca^2+^ imaged with synaptophysin-GCaMP6f in excitatory neurons with MCU knockdown (KD) and wild type (WT) during stimulation with 20 action potentials (AP), 20 Hz (n=9 WT, n=9 MCU KD, error bands SEM). **B)** Synaptic Ca^2+^ in inhibitory neurons with MCU KD in comparison with WT inhibitory neurons, during 20 AP, 20 Hz stimulation (n=8 WT, n=10 MCU KD, error bands SEM). **C)** Quantification of change in synaptic Ca^2+^ from A) and B). (n=9 excitatory WT, mean=99.2. n=9 excitatory MCU KD, mean=102.2. n=8 inhibitory WT, mean=75.01. n=10 inhibitory MCU KD, mean=193.8. Error bars SEM, n.s., p>0.99, **p=0.0014, Mann-Whitney test). **D)** Resting level of Ca^2+^ detected with physin-GCaMP6f in excitatory and inhibitory neurons with and without MCU KD (n=9 excitatory WT, mean=114.2. n=8 excitatory MCU KD, mean=114.6. n=8 inhibitory WT, mean=101.9. n=10 inhibitory MCU KD, mean=117.2. Error bars SEM, comparisons n.s., Kruskal-Wallis test). **E)** Synaptic vesicle cycling traces from excitatory neurons with MCU KD or WT controls (n=13 WT, n=15 MCU KD, error bands SEM). **F)** Synaptic vesicle cycling fluorescence traces from MCU KD and WT inhibitory neurons (n=10 WT, n=15 MCU KD, error bands SEM). **G)** Quantification of response amplitude from E) and F) (n=13 WT excitatory, mean= 0.07869. n=15 MCU KD excitatory, mean=0.0666. n=10 WT inhibitory, mean=0.0572. n=15 MCU KD inhibitory, mean=0.1327. Error bars SEM. **p=0.0044, n.s. p=0.9471, Mann-Whitney test). **H)** Proposed pathway of activity-dependent mitochondrial Ca^2+^ handling in inhibitory neurons.

### Inhibitory axonal mitochondria have faster Ca^2+^ efflux

Although mitochondrial Ca^2+^ uptake helps to accelerate oxidative phosphorylation, likely in part by modulating the activity of the matrix dehydrogenases (Denton, 2009; Gaspers & Thomas, 2008), excessive mitochondrial Ca^2+^ can be deleterious to cell health (Kruman & Mattson, 1999). Given that many inhibitory neurons have higher intrinsic firing rates, we wondered if the kinetics of mitochondrial Ca^2+^ extrusion are similar to those in excitatory neurons. We compared the dynamics of mitochondrial Ca^2+^ with Mito4x-jRCaMP1b in axons of excitatory and inhibitory neurons during stimulation. Remarkably, mitochondrial Ca^2+^ decay kinetics in inhibitory mitochondria were 6-fold faster in inhibitory neurons (Figure 4A, 4D; Supplementary Figure 3A, 3C). These results suggest a novel mechanism for differential regulation of mitochondrial metabolism by Ca^2+^ across neuronal subtypes.

**Figure 4.**
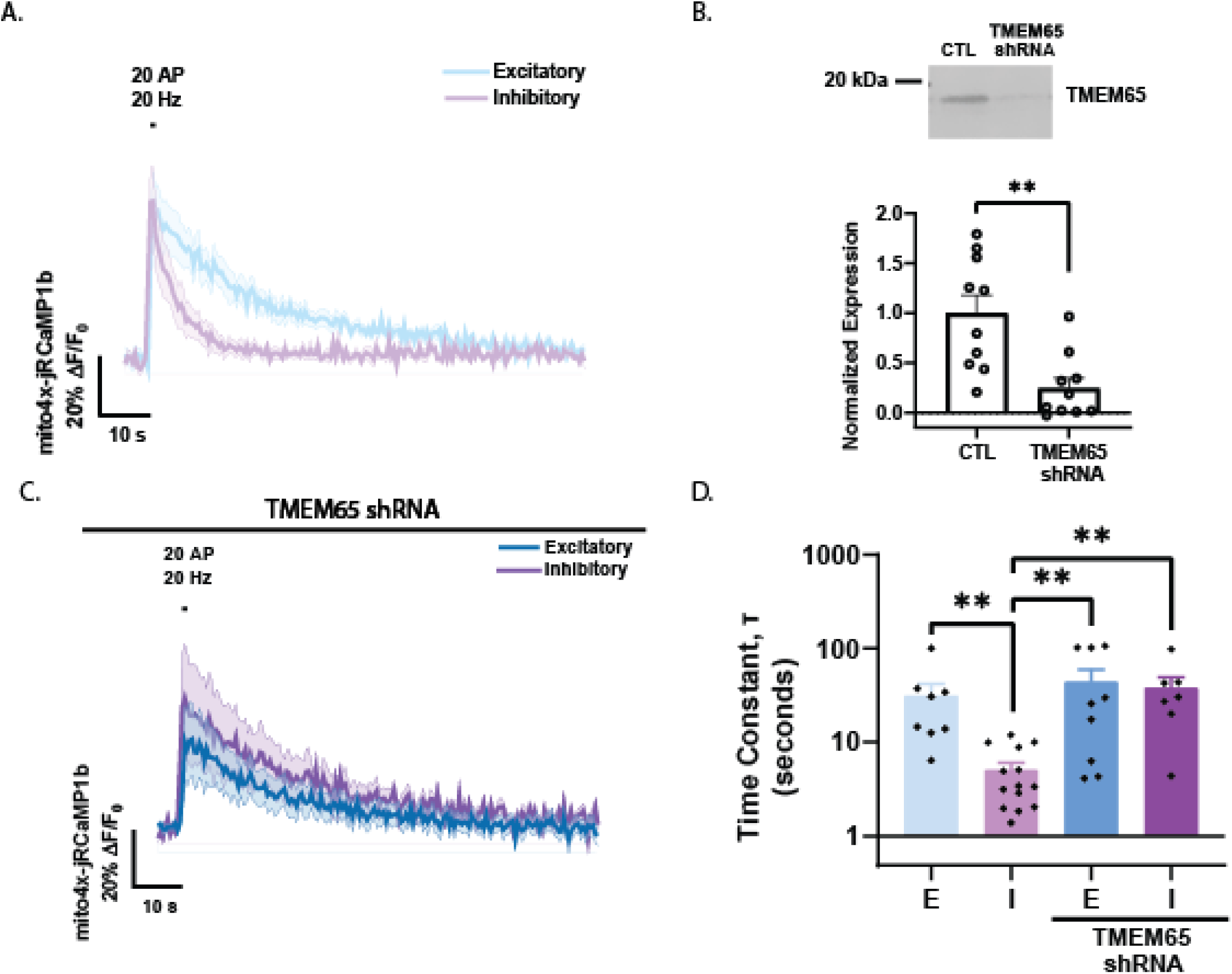
TMEM65 regulates mitochondrial Ca^2+^ efflux in inhibitory neurons. **A)** Mitochondrial Ca^2+^ detected with jRCaMP1b targeted to the mitochondrial matrix (Mito4x-jRCaMP1b) during 20 action potential (AP), 20 Hz stimulation (n=11 excitatory, 9 inhibitory, error bands SEM). **B)** (Above) Representative Western blot for TMEM65, comparing culture of cortical neurons treated with TMEM65 shRNA virus for 10 days and paired untreated control culture. (Below) Quantification of percent knockdown of TMEM65 by shRNA virus, detected by Western blot (N=5 cultures, n=2 technical replicates. TMEM65 shRNA mean 24.95% of CTL. Error bars SEM, **p=0.0021, Mann-Whitney test). **C)** Mitochondrial Ca^2+^ traces from excitatory and inhibitory neurons with shRNA mediated knock down of endogenous TMEM65 (n=8 excitatory, 9 inhibitory, error bands SEM). **D)** Quantification of time constants of Ca^2+^ extrusion (n=8 excitatory control, mean=31.38. n=14 inhibitory control, mean=5.055. n=9 excitatory TMEM65 shRNA, mean=44.28. n=7 inhibitory TMEM65 shRNA, mean=38.12. Error bars SEM, **p<0.01, Kruskal-Wallis test).

### TMEM65 mediates rapid Ca^2+^ efflux in inhibitory neurons

We reasoned that the observed higher rate of mitochondrial Ca^2+^ decay in inhibitory neurons most likely reflects a more efficient mitochondrial Ca^2+^ efflux pathway in this class of neurons. Mitochondrial Ca^2+^ is thought the be governed by a balance of the activity of the influx pathway (largely accounted for by MCU) and a mixture of extrusion mechanisms. Among these are the Ca^2+^/H^+^ exchanger (CHE) and the Na^+^/Ca^2+^ exchanger (NCLX) (Giorgi et al., 2018). Recently, TMEM65 was proposed as a putative regulator of NCLX that speeds Ca^2+^ efflux (Garbincius et al., 2023; Vetralla et al., 2023). Furthermore, TMEM65 mRNA is more abundant in inhibitory neurons than excitatory neurons (Huntley et al., 2020). Western blot analysis confirmed that TMEM65 is expressed in our primary cultured neurons (Figure 4B). To assess whether TMEM65 modulated Ca^2+^ efflux in neurons, we designed an shRNA targeting this protein that led to ∼ 80% knockdown in our system (Figure 4B). While expression of the TMEM65 shRNA had no impact on presynaptic cytoplasmic Ca^2+^ (either resting, or in response to activity, Supplementary Figure 4B-D), expression of the TMEM65 shRNA in inhibitory neurons significantly slowed mitochondrial Ca^2+^ extrusion following a burst of AP firing such that it was largely indistinguishable from the kinetics of mitochondrial Ca^2+^ extrusion in excitatory axons (Figure 4C, 4D). Expression of the shRNA in excitatory neurons in contrast had no measurable impact on the kinetics of mitochondrial Ca^2+^ extrusion (Figure 4C, 4D). These results therefore identify TMEM65 as a differential regulator of mitochondrial Ca^2+^ export in inhibitory neuron mitochondria.

Consistent with previous reports (Garbincius et al., 2023), loss of TMEM65 appeared to be detrimental to cell health, as further attempts to examine neuronal functioning using genetically encoded reporters revealed an inability to reliably observe synaptic responses. As we observed a higher resting mitochondrial Ca^2+^ concentration in neurons expressing TMEM65 shRNA (Supplementary Figure 4A), we attributed the decreased cell health to stress associated with mitochondrial Ca^2+^ overload.

## Discussion

Metabolism has often been considered an important contributor in controlling the balance between excitation and inhibition in the brain. Using a reductionist approach, we examined how excitatory and inhibitory neurons might differ in the metabolic control of axonal function. Our work revealed several important findings. We detected an ∼ 25% higher axonal mitochondrial density in inhibitory neurons, consistent with measurements in more intact tissues (Gulyás et al., 2006). This modest difference in mitochondrial density, however, had major ramifications with respect to bioenergetics. Under resting conditions, inhibitory neurons had almost double the concentration of ATP (Figure 1D). This difference appears to be entirely accounted for by the higher mitochondrial density in this neuron class, as blocking mitochondrial ATP production quickly eliminated the difference in resting ATP (Figure 1E), while switching to a purely respiratory fuel led to an even greater difference in ATP levels (Figure 2E). It is notable that this difference is apparent even in the absence of explicit electrical activity because it indicates that inhibitory neurons operate at a different set-point with regards to ATP levels. These results establish a key role for mitochondria in our understanding of E/I balance, particularly under conditions where alternative fuels may be supplied. For example, ketogenic diets have been shown to be effective in a large fraction of drug resistant childhood-epilepsies (Sourbron et al., 2020) and would be predicted to preferentially support inhibitory neuron function since this fuel relies purely on mitochondria for combustion and production of ATP. Lactate can similarly be exchanged across the plasma membrane via monocarboxylic acid transporters (MCTs), in principle allowing this glycolytic end-product to be used by other cells following conversion back to pyruvate (Roosterman & Cottrell, 2020). The difference in the reliance of excitatory versus inhibitory bioenergetics on mitochondrial function suggests that inhibitory neurons would be able to better make use of this indirect fueling pathway. Our results have strong implications with respect to the impact of lesions in the proper fueling of inhibitory neurons or the integrity of the mitochondrial machinery in these cells. We showed that the failure of mitochondrial Ca^2+^ uptake in inhibitory neurons leads to significant changes in the accumulation of Ca^2+^ in the presynaptic terminal that in turn increase rates of exocytosis, demonstrating that in inhibitory axons mitochondria play a significant role in Ca^2+^ buffering. Presumably, if mitochondria failed to maintain their membrane potential, then the metabolic burden in synapses would increase because of increased exocytosis, further taxing the metabolic capacity of the neuron where mitochondria are not running at full efficiency.

It is notable that despite the greater respiratory capacity in inhibitory axons, the dynamics of ATP changes during intense electrical stimulation in glucose were similar to those seen in excitatory axons (Figure 2A). Both the activity driven ATP deficit (Figure 2B) and recovery following stimulation (Supplementary Figure 1B) were similar in the two neuron classes, suggesting that acute ATP dynamics are governed largely by glycolysis, consistent with our recent discovery that the first ATP producing enzyme in glycolysis (phospoglycerate kinase 1) is rate limiting (Kokotos et al., 2023). The lack of a difference in excitatory and inhibitory ATP recovery rates following stimulation even in conditions of metabolic compromise (Supplementary Figure 1C, 1D, 1F) suggests that ATP consumption and production rates during and directly following stimulation are similarly well equipped to meet metabolic need in excitatory and inhibitory neurons. The difference in baseline ATP levels between excitatory and inhibitory axons in oxidative conditions (Figure 2E) but not in axons operating under anaerobic glycolysis (Figure 1E) suggests that inhibitory axons may rely on unique Ca^2+^ dependent mechanisms of upregulating mitochondrial metabolism to maintain their higher ATP levels. The function of Ca^2+^ in neuronal mitochondria is undoubtedly complex but still incompletely understood (Giorgi et al., 2018). Others have suggested a role for mitochondrial Ca^2+^ buffering in uncoupling mitochondrial ATP synthesis (Werth & Thayer, 1994), such that mitochondrial Ca^2+^ export is essential to mitochondrial ATP production through restoration of the mitochondrial membrane potential, and perhaps even more important than the role of Ca^2+^ in activating mitochondrial dehydrogenases to meet activity-related metabolic needs with mitochondrial metabolism. If so, fast Ca^2+^ export by TMEM65 may prevent mitochondrial Ca^2+^ accumulation in highly active neurons from reducing mitochondrial membrane potential.

Whether the mitochondrial dehydrogenases are activated by Ca^2+^ in a tonic or phasic manner remains largely unknown, and therefore whether Ca^2+^ efflux mediated by TMEM65 would affect metabolic activation. According to recent work, pyruvate dehydrogenase regulation by Ca^2+^ occurs indirectly through calmodulin-dependent activation of pyruvate dehydrogenase phosphatase that persists long enough to encode neuronal activity level (Yang et al., 2024). Since calmodulin cooperatively binds four Ca^2+^ ions, greater concentrations of Ca^2+^ might lead to a longer lasting conformational change, although the consequences of these changes for binding to target proteins are not well understood (Stigler & Rief, 2012). While this mechanism suggests phasic activation, the TCA cycle dehydrogenases may still be activated tonically. Oxoglutarate and isocitrate dehydrogenases (OGDH and IDH, respectively) bind Ca^2+^ directly at largely undescribed sites. The affinity of OGDH for Ca^2+^ alsodepends on the ATP/ADP ratio (Wan et al., 1989; Denton, 2009), further adding to the complexity of activity-dependent regulation. In sum, a slower efflux of Ca^2+^ in excitatory neurons could hypothetically keep the TCA cycle active longer than in inhibitory neurons if OGDH and IDH are activated tonically, but more work is required to understand the function of changing mitochondrial Ca^2+^ kinetics.

In summary, we have found that inhibitory neurons utilize oxidative phosphorylation more than excitatory neurons when fueling neurotransmission and rely in part on differential expression of a putative NCLX regulator, TMEM65, to accelerate mitochondrial Ca^2+^ extrusion following electrical activity. Our studies suggest that many other key molecular differences may exist between inhibitory and excitatory mitochondria, a concept consistent with transcriptomics of mitochondrial genes in different cell types (Wynne et al., 2021). Purification of mitochondria from different neuron classes (Fecher et al., 2019) should prove fruitful in future approaches.

## Acknowledgements

This work was supported by NIH grant 2R01NS036942 awarded to T.A.R. and by a Medical Scientist Training Program grant from the National Institute of General Medical Sciences of the NIH under award number T32GM007739 to the Weill Cornell/Rockefeller/Sloan Kettering Tri-Institutional MD-PhD Program. We thank Giulia Dalaty and Hayoung Lee for technical assistance, and we thank members of the Ryan lab for helpful discussion, especially Alexandros Kokotos for generating the pLKO.1 hPGK-BFP TRC cloning vector, and Ryan Farrell, Ghazaleh Ashrafi and Jaime de Juan-Sanz for essential advice and for generating the Mito4x-jRCaMP1b construct. Some figures were generated using Biorender.

## Author Contributions

K.B. and T.A.R. contributed to the experimental design. K.B. collected and analyzed data.

K.B. and T.A.R. contributed to writing the manuscript.

## Materials and Methods

### Animals

All experiments were performed using 1-to 2-day old wild type Sprague-Dawley rats (Charles River code 400, RRID: RGD_734476). Rats of mixed gender were sacrificed and dissected according to protocols approved by Weill Cornell Medicine IACUC.

### Culture and Transfection

Neurons were dissected from the CA1 to CA3 regions of the hippocampus, or from the cortex and hippocampus of 1- to 3-day-old rats. Neurons were cultured and plated on coverslips coated with poly-L-ornithine as previously described (Farrell et al., 2023). Calcium phosphate-mediated gene transfer over a 1 h incubation was used to transfect 6- to 7-day-old cultures after three washes with DMEM, as described previously (Sankaranarayanan et al., 2000). Neurons were incubated at 37° C in a 95% air/5% CO_2_ humidified incubator until imaged 14-21 days after plating. Culture media consisted of MEM (Thermo Fisher #51200038), 0.25 g/L insulin, 0.3 g/L GlutaMAX, 5% fetal bovine serum (Atlanta Biological S11510), 2% N-21 (Chen et al., 2008), and 4 µM cytosine β-d-arabinofuranoside.

#### Western blots

Cortical neurons were cultured from 1- to 3-day-old rats and plated in 6-well plates coated with poly-L-ornithine. Cytosine β-d-arabinofuranoside was added to the media after 2-3 days, using the same media formulations as for hippocampal cultures (Farrell et al., 2023).

Approximately 500,000 GC of lentivirus was added to each well containing a mixture of neurons and glia totaling about 2.5 million cells. Cultures were harvested after 10-11 days incubation with lentivirus and samples were prepared for western blot using RIPA buffer (Thermo Fisher, 89900). Samples were heated for 10 minutes at 95° C with 50 mM DL-dithiothreitol (Sigma Aldrich, 43819) and Laemmli buffer at 1X (Bio-Rad, 161-0747) before loading into 4-20% precast gels (Bio-Rad, 4561096 and 4561094). After running for 45 minutes at 200 V in Tris/Glycine/SDS buffer (Bio-Rad, 1610732), gel was transferred to nitrocellulose membrane (Bio-Rad, #1620115) at 100 V for 30 minutes at 4° C in Tris/Glycine buffer (Bio-Rad, 1610734) with 20% methanol. Protein was quantified using REVERT™ total protein stain (LI-COR, #926-11011) per manufacturer instructions. Membranes were blocked for 1 h with Odyssey Blocking Buffer (TBS) (LI-COR, #927-50000) and incubated overnight at 4° C in primary antibody and Odyssey Blocking Buffer. TMEM65 rabbit polyclonal antibody (Proteintech 21913-1-AP) was used at 1:10,000. Secondary antibody (IRDye^®^ 800CW Goat Anti-Rabbit, LI-COR #926-32211) was incubated for 1 h at room temperature, diluted 1:20,000 in Odyssey Blocking Buffer after washing with TBS + 0.1% Tween-20. LI-COR Odyssey^®^ Fc Imaging System and Image Studio Software was used to develop and quantify blots.

#### Analysis

Western blots were analyzed using the “draw rectangles” function in LI-COR Image Studio Lite software. The signal for each lane was normalized by total protein signal (REVERT™ total protein stain). For graphs, each replicate was additionally normalized to the control average.

Images were analyzed with the Time Series Analyzer plugin in ImageJ, using 20-149 ROIs of ∼ 2 μm diameter to measure the fluorescence of synaptic boutons over time. To quantify change in fluorescence at a specific time point, the fluorescence of three frames (pHluorin) or five frames (iATPSnFR2.0-HALO) were averaged and compared to a pre-stimulation baseline of between 10 and 25 frames. iATPSnFR2.0-HALO measurements were taken using alternating frames of 488 and 637 laser illumination, where the cell average of regions from each frame imaged with 488 laser was normalized to the paired cell average of 10 frames imaged with 637 laser prior to stimulation.

### Imaging

Live imaging experiments were performed on a custom-built laser illuminated epifluorescence microscope with an Andor iXon+ camera (model DU-897E-CS0-#BV). Coherent OBIS 488 nm, 561 nm, and 637 nm lasers illuminated samples under control of an Arduino program. Images were acquired through a 40X 1.3 NA Fluar Zeiss objective. Coverslips were mounted in a laminar flow perfusion chamber and perfused with Tyrode’s buffer containing (in mM) 119 NaCl, 2.5 KCl, 2 CaCl_2_, 2 MgCl_2_, 50 HEPES (pH 7.4), 5 glucose or 1.25 lactate and 1.25 pyruvate, supplemented with 10 μM 6-cyano-7-nitroquinoxalibe-2, 3-dione (CNQX), and 50 μM D,L-2-amino-5phosphonovaleric acid (APV) (both from Sigma-Aldrich) to prevent excitatory post-synaptic responses. Tyrode’s buffer containing 1.1 mM glucose was supplemented with oligomycin (Sigma Aldrich) at a concentration of 6 µM when specified. Tyrode’s with lactate/pyruvate, oligomycin, or CPA (Alomone) were perfused for 5 minutes prior to imaging to allow for complete replacement of fluid in perfusion chamber. Action potentials were evoked in neurons with 1 ms pulses between platinum–iridium electrodes resulting in a field potential of ∼10 V/cm. Stimulation timing was controlled by custom-built Arduino and python programs. Temperature was maintained at 37°C using a custom-built objective heating jacket.

Prior to iATPSnFR2.0 imaging, cells were incubated with 50 nM Janelia Fluor® 635 HaloTag® ligand (JF635) dye for 30 minutes, then washed with Tyrode’s buffer. For some iATPSnFR2.0 imaging, cells were initially transfected with CAG iATPSnFR2.0-HALO and each dish was treated with 1:100 VGAT Oyster 550 lumenal antibody (Synaptic Systems) during 600 AP, 10 Hz stimulation to identify inhibitory neurons.

Mitochondrial and synaptic resting free Ca^2+^ concentrations for cells expressing Mito4x-jRCaMP1b or synaptophysin-GCaMP6f were calculated by measuring the fluorescence at saturating [Ca^2+^], F_max_, by applying ionomycin (Alomone Labs) at 500 μM in Tyrode’s buffer containing (in mM) 119 mM NaCl, 2.5 KCl, 4 CaCl_2_, 55 HEPES (pH 6.9). The baseline Ca^2+^ was then calculated using the known in vitro characteristics of GCaMP6f (Chen et al., 2013) and jRCaMP1b (Dana et al., 2016) using the following equation:

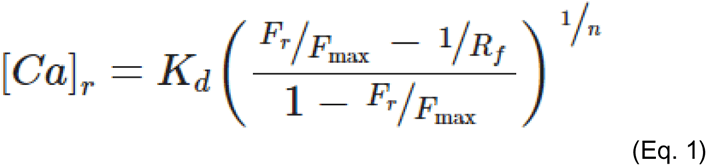

NH_4_Cl solution used for pHluorin-containing vesicle alkalization contained (in mM) 50 NH_4_Cl, 69 mM NaCl, 2.5 KCl, 2 CaCl_2_, 2 MgCl_2_, 55 HEPES (pH 7.4).

#### Plasmid Constructs

CaMKII synaptophysin-pHluorin was made by digestion of CaMKII ER-GCaMP3-373 (de Juan-Sanz et al., 2017) and joined by Gibson HiFi reaction with PCR of synaptophysin-pHluorin (Pan et al., 2015). To make hDlxI56i synaptophysin-pHluorin, synaptophysin-pHluorin was cut with NheI and MluI, and joined by Gibson HiFi reaction (NEB) with PCR of the hDlxI56i enhancer and minimal beta globin promoter from hDlxI56i-minBglobin-iCre-4X2C (Addgene #164450). hDlxI56i eGFP plasmid was made by cutting CMV-eGFP-synapsin (Chi et al., 2001) with AgeI, AseI, BamHI, and BglII, and performing PCR of hDlxI56i from hDlxI56i synaptophysin-pHluorin, ligating with Quick Ligase and performing Gibson HiFi reaction.

To make hDlxI56i synaptophysin-GCaMP6f, hDlxI56i synaptophysin-pHluorin was cut with AflII and XhoI, and Gibson HiFi reaction was used to add synaptophysin-GCaMP6f from PCR of hSyn synaptophysin-GCaMP6f (de Juan-Sanz et al., 2017). To make CaMKII synaptophysin-GCaMP6f, CaMKII-ER-GCaMP6f (de Juan-Sanz et al., 2017) was cut with BamHI and EcoRI and Gibson HiFi reaction was used to join with PCR from synaptophysin-GCaMP6f. CaMKII Mito4x-jRCaMP1b and hDlxI56i Mito4x-jRCaMP1b were made by cutting CMV-Mito4x-jRCaMP1b (Addgene #127873) with MluI and BamHI and performing PCR of CaMKII and hDlxI56i from CaMKII and hDlxI56i synaptophysin-pHluorin, respectively.

hDlxI56i iATPSnFR2.0-HALO was made by digesting hDlxI56i eGFP with MfeI and AgeI, and using Gibson HiFi to combine with PCR from CaMKII iATPSnFR2.0-HALO (Marvin et al., 2024). All shRNA plasmids were made by digesting pLKO.1 hPGK-BFP TRC cloning vector (Addgene #191566) with AgeI-HF and EcoRI-HF and ligating annealed shRNA sequences with Quick Ligase Buffer (NEB, M2200). Primers for oligos against the listed sequences followed the format:

5’ CCGG-sense-CTCGAG-antisense-TTTTTG 3’, 5’AATTCAAAAA-sense-CTCGAG-antisense 3’

Target sequences:

MCU: GCTACCTTCTCGGCGAGAACGCTGCCAGTT

TMEM65: ATTGTTGCAGGAACCCAAATT

#### Virus production

HEK293FT cells were transfected by calcium phosphate with lentiviral constructs along with the associated packaging plasmids psPAX2 (a gift of Didier Trono, Addgene #12260) and pMD2.G (a gift of Didier Trono, Addgene 12259). 16 h post transfection, media was changed to serum free viral production media: Ultraculture (Lonza), 1% (v/v) Penicillin-Streptomycin/L-glutamine, 1% (v/v) 100 mM sodium pyruvate, 1% (v/v) 7.5% sodium bicarbonate and 5 mM sodium butyrate. HEK293FT supernatants were collected at 46 h post transfection and filtered through a 0.45 mm cellulose acetate filter. Viral supernatants were then concentrated using Lenti-X Concentrator (Takara) and spun for 45 min at 1500 x g before resuspending in PBS. Samples were aliquoted and stored at 80°C until use. Genomic titer was determined by Lenti-X GoStix Plus (Takara). Lentivirus was functionally titrated previously using parallel preparation of GFP-expressing viral particles (Campeau et al., 2009) to determine the volume of lentivirus required to achieve ∼ 100% transduction of hippocampal or cortical cultures, 106 GC (genome copies)/mL (Ashrafi et al., 2020). Lentivirus was added to neurons 2-4 days *in vitro*, and all experiments were performed at least 10 days after viral transduction to ensure protein knockdown.

### Statistical Analysis

Fitting was done in GraphPad Prism v8, using the exponential decay function without constraints. For endocytic rates of fluorescence decay, fits were made using data points starting 2 seconds after the end of stimulation. For mitochondrial Ca^2+^ traces, fits included data points starting at the end of stimulation. Statistical analysis was performed with GraphPad Prism v8. As specified in the figure legends, the significance of differences between conditions was calculated using the nonparametric Mann-Whitney test for unpaired or the Kruskal-Wallis test for paired comparisons, because we did not assume that the distributions of our datasets would follow a normal distribution. Dunn’s correction was used to correct for multiple comparisons. Generally, p < 0.05 was considered significant and denoted with one asterisk, while p < 0.01, p < 0.001, p < 0.0001 were shown with two, three, or four asterisks respectively.

### Supplemental material

Supplement includes figures and figure legends with expanded quantifications of data from main figures, as well as additional experiments that clarify interpretation of main figure data.

