## Supplemental Figures for "Differential Control of Inhibitory and Excitatory Nerve Terminal Function by Mitochondria"

### Supplementary Figure 1.

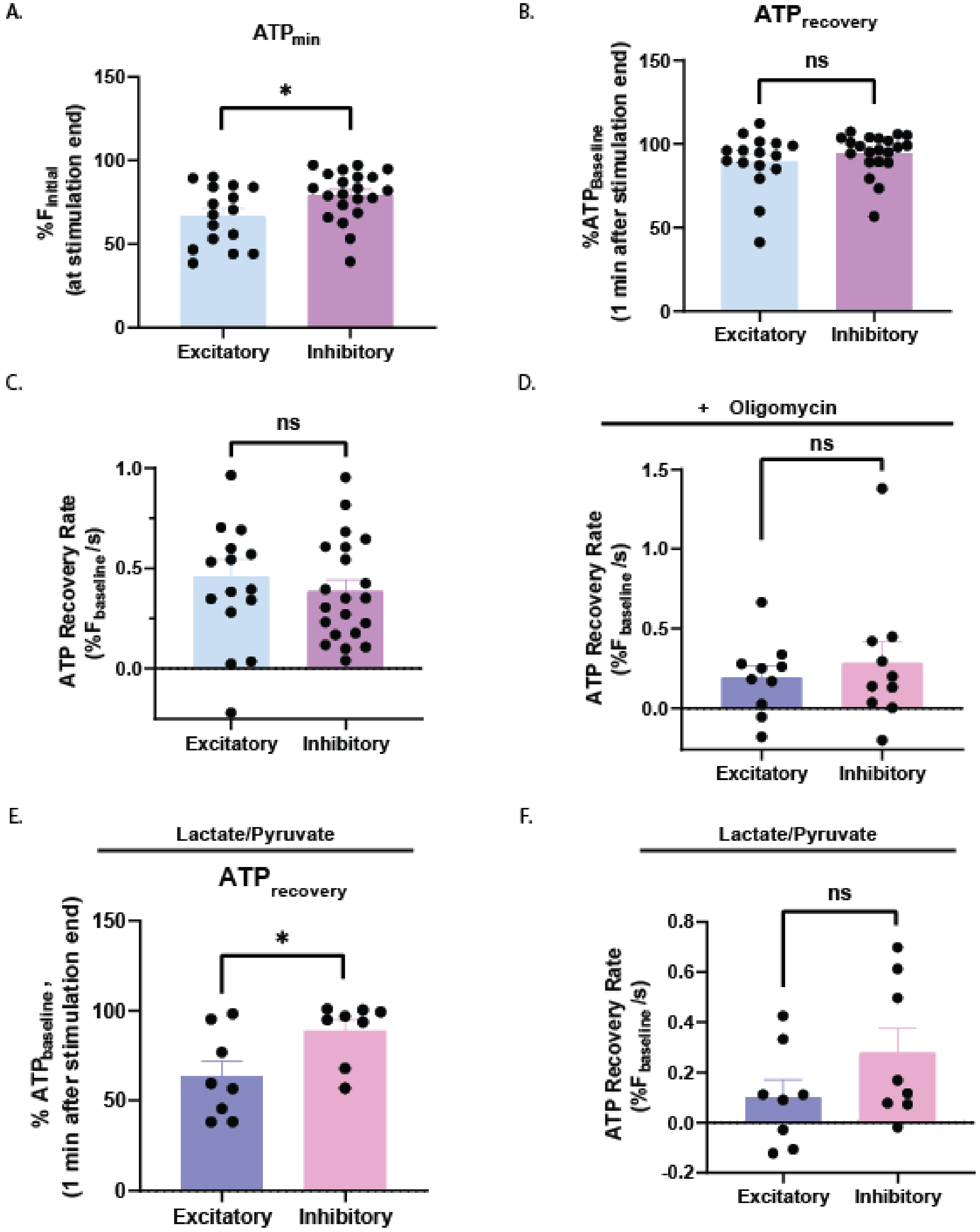

**Supplementary Figure 2.**

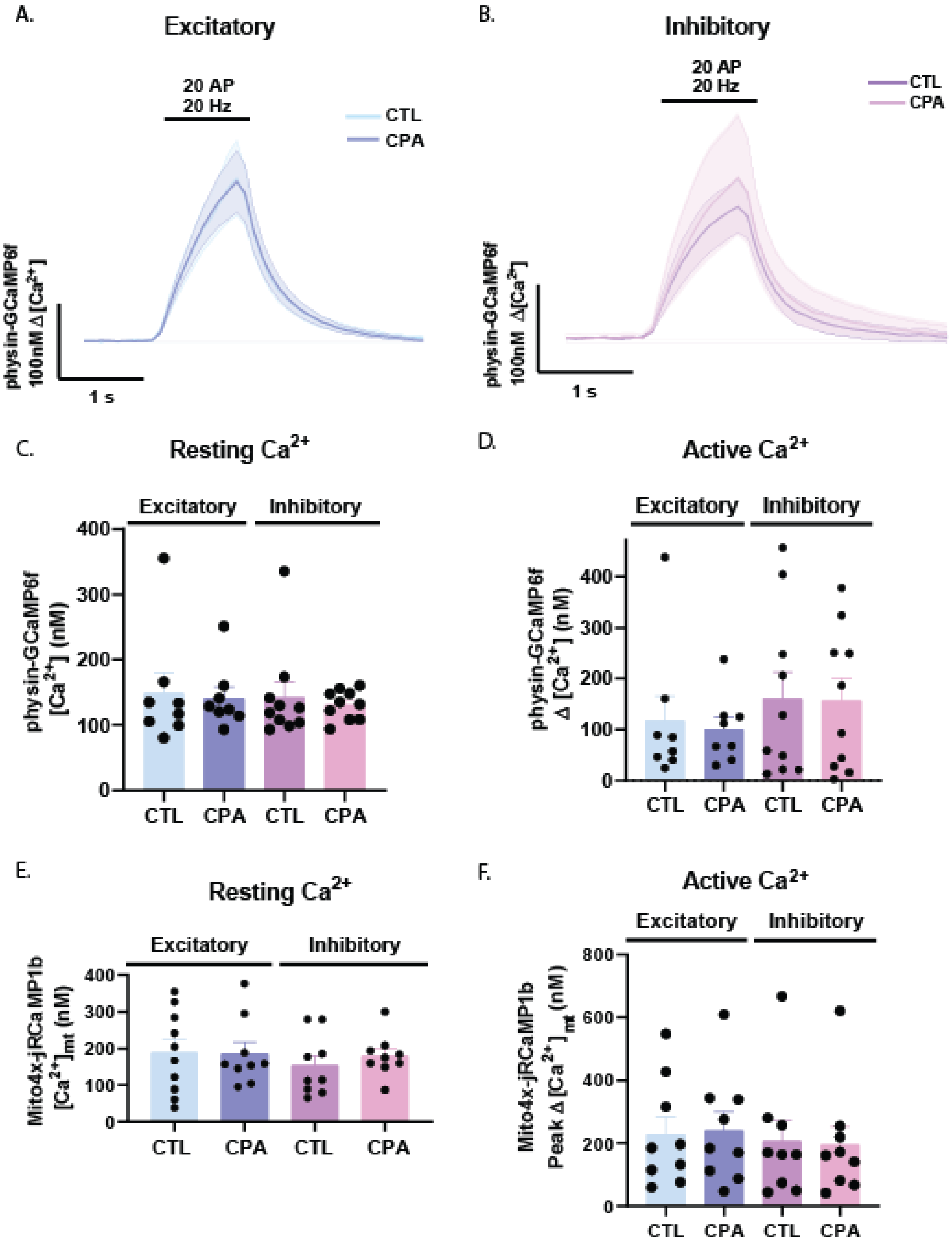

**Supplementary Figure 3.**

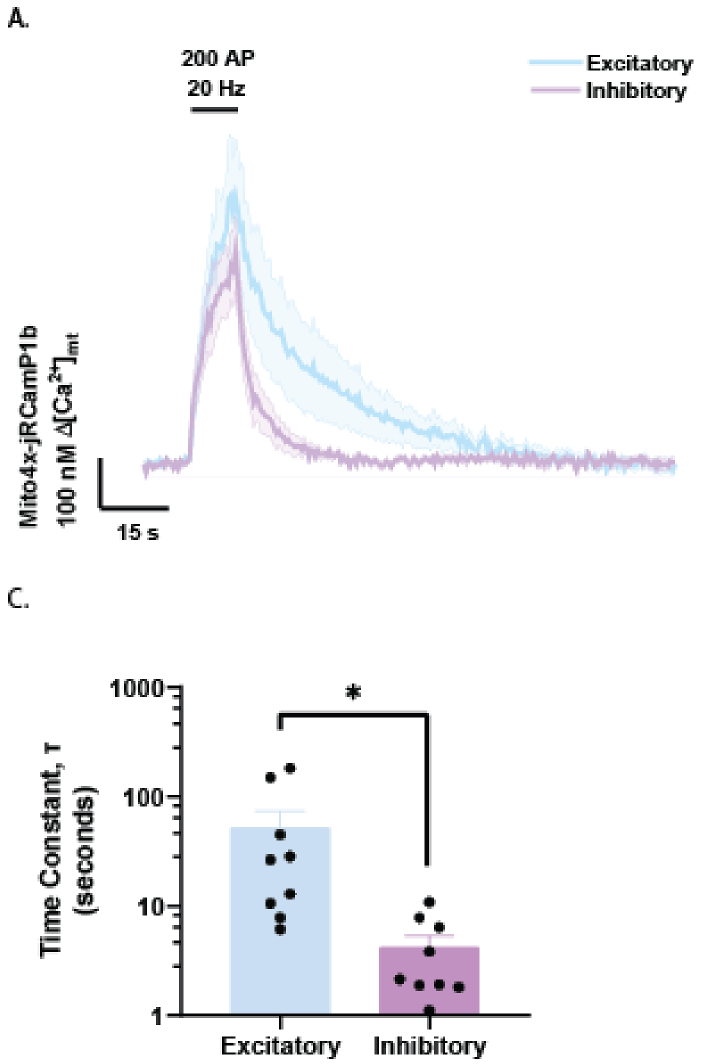

**Supplementary Figure 4.**

A.

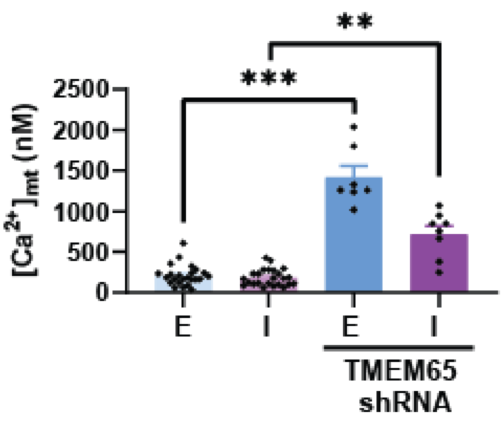


B.

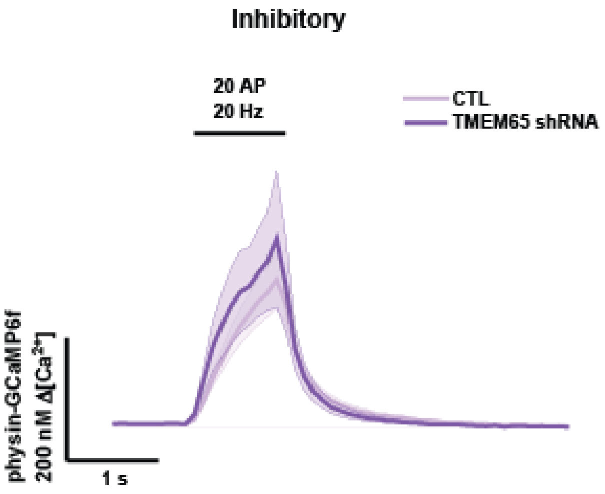


C.

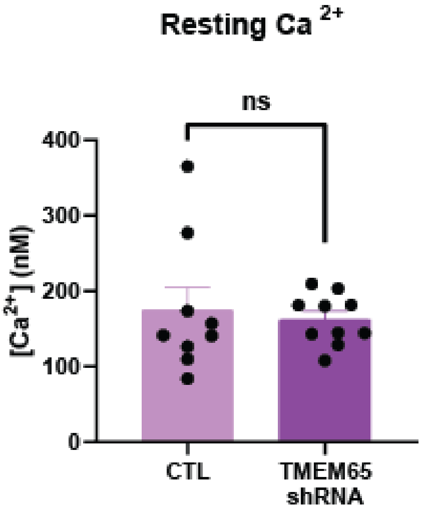


D.

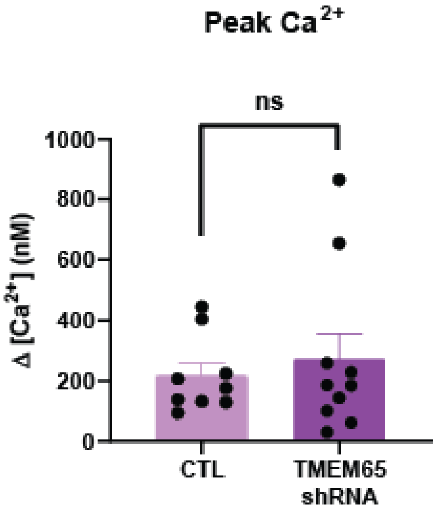

### Supplementary Figure Legends

**Figure S1. Excitatory and inhibitory ATP levels and synaptic transmission are similarly affected by oligomycin treatment. A)** Quantification of ATP following stimulation as percentage of baseline, from traces in Figure 2A (n=16 excitatory, mean=66.7%, n=21 inhibitory, mean=79.49%. Error bars SEM, *p=0.0346, Mann-Whitney test). **B)** Quantification of ATP one minute after cessation of 600 AP, 10 Hz stimulation (n=16 excitatory, mean=89.31%, n=21 inhibitory, mean=94.43%, error bars SEM. n.s., p=0.2537, Mann-Whitney test). **C)** Rate of ATP recovery quantified as percent of baseline fluorescence per second after 600 action potential (AP), 10 Hz stimulation, from traces in Figure 2A (n=16 excitatory, mean=0.4978. n=21 inhibitory, mean=0.3873. Error bars SEM, n.s., p=0.2941, Mann-Whitney test). **D)** Rate of ATP recovery following 600 AP, 10 Hz stimulation with oligomycin treatment, from oligomycin traces in Figure 2C (n=10, excitatory mean=0.194, inhibitory mean=0.2856. Error bars SEM, n.s., p=0.9118, Mann-Whitney test). **E)** Quantification of normalized fluorescence ratio one minute after cessation of 600 AP, 10 Hz stimulation in 1.25 mM lactate and 1.25 mM pyruvate, from traces in Figure 2F (n=8, excitatory mean=63.63%, inhibitory mean=88.9%. Error bars SEM, *p=0.0379, Mann-Whitney test). **F)** Quantification of rate of recovery as percent of baseline fluorescence per second, following stimulation of 600 AP, 10 Hz in lactate/pyruvate, from traces in Figure 2F (n=8, excitatory mean=0.1019, inhibitory mean=0.2784. Error bars SEM, n.s., p=0.1949, Mann-Whitney test).

**Figure S2. Endoplasmic reticulum $Ca^{2+}$ stores are not major contributors to mitochondrial calcium entry during activity.** Average synaptic $Ca^{2+}$ uptake detected with synaptophysin-GCaMP6f (physin-GCaMP6f) in response to a 20 action potential (AP), 20 Hz stimulation before and after 5 minute cyclopiazonic acid (CPA) treatment in **A)** excitatory neurons (n=8, error bands SEM) and **B)** inhibitory neurons (n=10, error bands SEM). **C)** Average resting synaptic $Ca^{2+}$ in excitatory and inhibitory neurons before and after CPA treatment (n=8 excitatory, 10 inhibitory. Excitatory CTL mean=149.2, excitatory CPA mean=141.6, inhibitory CTL mean=143, inhibitory CPA mean= 131.3, error bars SEM). **D)** Quantification of change in synaptic $Ca^{2+}$ with 20 AP, 20 Hz stimulation from A) and B) (n=8 excitatory, n=10 inhibitory, excitatory CTL mean=117.9, excitatory CPA mean=101.1, inhibitory CTL mean=160.9, inhibitory CPA mean=157.3, error bars SEM). **E)** Quantification of resting mitochondrial $Ca^{2+}$ detected with Mito4x-jRCaMP1b, before and after CPA treatment (n=10 excitatory CTL, mean=188.9, n=9 excitatory CPA, mean=186, n=9 inhibitory CTL, mean=153.5, n=9 inhibitory CPA, mean=179.5, error bars SEM). **F)** Quantification of peak mitochondrial $Ca^{2+}$ entry following stimulation with 20 AP, 20 Hz (n=9 excitatory, n=9 inhibitory, excitatory CTL mean=228.2, excitatory CPA mean=240.9, inhibitory CTL mean=207.5, inhibitory CPA mean=195.1, error bars SEM).

**Figure S3. $[Ca^{2+}]_{mt}$ decay in strongly stimulated inhibitory neurons is faster than in excitatory neurons. A)** Change in mitochondrial $Ca^{2+}$ in response to 200 AP, 20 Hz stimulation of neurons transfected with Mito4x-jRCaMP1b using excitatory and inhibitory neuron specific constructs (n=7 excitatory, n=6 inhibitory, error bands SEM). **B)** Quantification of peak change in mitochondrial $Ca^{2+}$ signal from A) (Excitatory mean=146.6, inhibitory mean=234.2. Error bars SEM, n.s, p=0.9452, Mann-Whitney test). **C)** Quantification of time constant of fluorescence decay after 200 AP, 20 Hz stimulation from A) (Excitatory mean=51.83, inhibitory mean=4.174. Error bars SEM, *p=0.0313, Mann- Whitney test).

**Figure S4. TMEM65 knockdown does not impact synaptic $Ca^{2+}$ dynamics of inhibitory synapses. A)** Resting mitochondrial $Ca^{2+}$ in neurons with TMEM65 shRNA and controls (n= 24 excitatory WT, mean=210.3, n=24 inhibitory WT, mean=184.3, n=7 excitatory TMEM65 sh, mean= 1419. n=8 inhibitory TMEM65 sh, mean=721.1. Error bars SEM, **p=0.0017, ***p=0.0001, Kruskal-Wallis test for multiple comparisons). **B)** Synaptic $Ca^{2+}$ detected by synaptophysin-GCamP6f (physin-GCaMP6f) during 20 action potential (AP), 20 Hz stimulation of inhibitory neurons with TMEM65 shRNA compared to control (n=9 control, n=10 TMEM65 shRNA, error bands SEM). **C)** Quantification of baseline synaptic $Ca^{2+}$ level prior to stimulation, from traces in A) (n=9 control, mean=170.7, n=10 TMEM65 shRNA, mean=156.6, error bars SEM. n.s., p=0.6047, Mann-Whitney test). **D)** Quantification of increase in synaptic $Ca^{2+}$ concentration during 20 AP, 20 Hz stimulation, from traces in A) (CTL mean=134.9, TMEM65 shRNA mean=255.8, error bars SEM. n.s., p=0.9048, Mann-Whitney test).
